# Synchrotron-source micro-x-ray computed tomography for examining butterfly eyes

**DOI:** 10.1101/2023.12.08.570583

**Authors:** Dawn Paukner, Gregg A. Wildenberg, Griffin S. Badalamente, Peter B. Littlewood, Marcus R. Kronforst, Stephanie E. Palmer, Narayanan Kasthuri

**Affiliations:** University of Chicago, Department of Neurobiology; Argonne National Laboratory; University of Cambridge, Department of Zoology; University of Chicago, Department of Physics; University of Chicago, Department of Organismal Biology and Anatomy; University of Chicago, Department of Ecology and Evolution University of Chicago: 5801 S Ellis Ave, Chicago, IL 60637 Argonne National Laboratory: 9700 S Cass Ave, Lemont, IL 60439 University of Cambridge: The Old Schools, Trinity Ln, Cambridge CB2 1TN, UK

**Author notes:** <u>Corresponding Author</u> Dawn Paukner 947-51 E. 58th St., AB 401 Chicago, IL 60637.

## Abstract

1. Comparative anatomy is an important tool for investigating evolutionary relationships amongst species, but the lack of scalable imaging tools and stains for rapidly mapping the microscale anatomies of related species poses a major impediment to using comparative anatomy approaches for identifying evolutionary adaptations.

2. We describe a method using synchrotron source micro-x-ray computed tomography (syn-µXCT) combined with machine learning algorithms for high-throughput imaging of Lepidoptera (*i.e.,* butterfly and moth) eyes. Our pipeline allows for imaging at rates of ∼ 15 min/mm^3^ at 600 nm^3^ resolution. Image contrast is generated using standard electron microscopy labeling approaches (e.g., osmium tetroxide) that unbiasedly labels all cellular membranes in a species independent manner thus removing any barrier to imaging any species of interest.

3. To demonstrate the power of the method, we analyzed the 3D morphologies of butterfly crystalline cones, a part of the visual system associated with acuity and sensitivity and found significant variation within six butterfly individuals. Despite this variation, a classic measure of optimization, the ratio of interommatidial angle to resolving power of ommatidia, largely agrees with early work on eye geometry across species.

4. We show that this method can successfully be used to determine compound eye organization and crystalline cone morphology. Our novel pipeline provides for fast, scalable visualization and analysis of eye anatomies that can be applied to any arthropod species, enabling new questions about evolutionary adaptations of compound eyes and beyond.

## Introduction

There is a rich history of using insects to understand behavioral and anatomical diversity (Chown & Terblanche, 2006; Price et al., 2011). Insects represent the largest group in the animal kingdom and their absolute numbers are also matched by their diversity in phenotypes, behavior, and anatomy (Stork, 2017). Classically, morphological variation that could be observed by the naked eye provided the necessary evidence for fundamental theories in evolution including natural selection, speciation, mimicry, and mate preference (Darwin, 1859; Poulton, 1909; Butler, 1963), to name a few. More recently, the revolution in genetics and genomics has allowed for identifying genetic variation that drives variation in these observable traits (Dobzhansky, 1982; Kronforst et al., 2006; Baxter et al. 2010). However, microscopic studies have lagged behind, largely due to a lack of experimental tools to rapidly visualize and analyze fine structural detail over large volumes and algorithmic tools to analyze the resulting large image data sets with minimal human effort. While there has been a recent push to test different techniques for studying morphology, most methods do not provide a satisfactory balance between higher resolution and lower computational power (Friedrich et al., 2014; Wipfler et al., 2016).

Electron microscopy (EM) can provide the requisite resolution but is typically limited to scanned EM, (SEM) which visualizes external morphologies (Schwarz et al., 2011; Hao et al., 2023). A full 3D EM reconstruction using serial block face SEM, focused ion beam SEM, or transmission EM remains time- and computation-intensive. We, and others, have recently shown that the sample preparation for EM using osmium tetroxide, which is species independent, provides excellent contrast in X-ray tomography microscopes (Johnson et al, 2006; Ribi et al, 2008; Dyer et al, 2017; Van den Boogert et al, 2018). Using X-ray tomography, large volumes of brains (even entire mouse brains) can be imaged in 3D at submicron resolution quickly (imaging rates of 0.067 mm^3^/min) (Foxley et al, 2021). Here we demonstrate a pipeline for synchrotron source X-ray computed tomography (syn-µCT) performed at the Advanced Photon Source (APS) at Argonne National Laboratory (ANL) for high throughput 3D imaging of the brains and intact eyes of a variety of butterflies.

1. We achieve 600nm^3^ voxel resolution and imaging rates of 0.067 mm^3^/min, e.g., ∼one insect brain every ∼45 minutes.
2. We developed a novel embedding method that allows for automatically imaging multiple species eyes in a single imaging run to enable high-throughput imaging.
3. We developed a machine vision pipeline to extract the relevant morphological features from X-ray datasets and used these reconstructions to better understand microscopic variability in the morphology of cells in the light path across species.
4. Specifically, we analyze these new data sets in the context of pioneering work in Hymenoptera species (e.g., bees and parasitic wasps) that determined an optimal ratio of interommatidial angle to resolving power (Barlow, 1952). This ratio of interommatidial angle to resolving power is hereafter referred to as the “Barlow ratio” and is dimensionless as both angle and resolving power are in degrees. We extend this work by showing the Barlow ratio of the ommatidia in butterfly species falls near the theoretical optimum. By leveraging the full 3D datasets, we, however, find significant variation *across an individual* eye.
5. Finally, we use an amalgamation of individual crystalline cone measurements across individual eyes to generate a representative 3D crystalline cone for each sample within and across species. Generating the morphology of these cones allows for the mapping of light as it travels through this structure to the rhabdom. We observe cone shapes that vary both across the eye of an individual and between individuals (Fig. 5). This technique allows for the dissection of these effects at fine detail across the eye and could support studies of cone optics and variation in and between species.

## Materials and Methods

Samples from seven animals across six species of butterflies (*Heliconius cydno*, *Strymon melinus*, *Calycopis cecrops*, *Polygonia interrogationis*, *Polites peckius*, and two *Pieris rapae*) were prepared for electron microscopy (Hua et al., 2015) and assembled in plastic pillars vertically to stabilize the samples for imaging (Fig. 1A), and large sections of eyes were imaged at the Advanced Photon Source (APS) at Argonne National Laboratory using syn-µCT) using an automated z-axis tiling approach for unassisted imaging of multiple insect eyes (Fig. 1B). The resulting X-ray data sets, with a total volume of 14.3 mm^3^ and an isotropic resolution of ∼0.6 microns resolved fine structure in the eye across all species, most notably the crystalline cones (Fig. 1C, Fig. SI1). We next developed our analysis pipeline by focusing on the crystalline cones due to its notable variability across species upon visual inspection.

**Figure 1.**
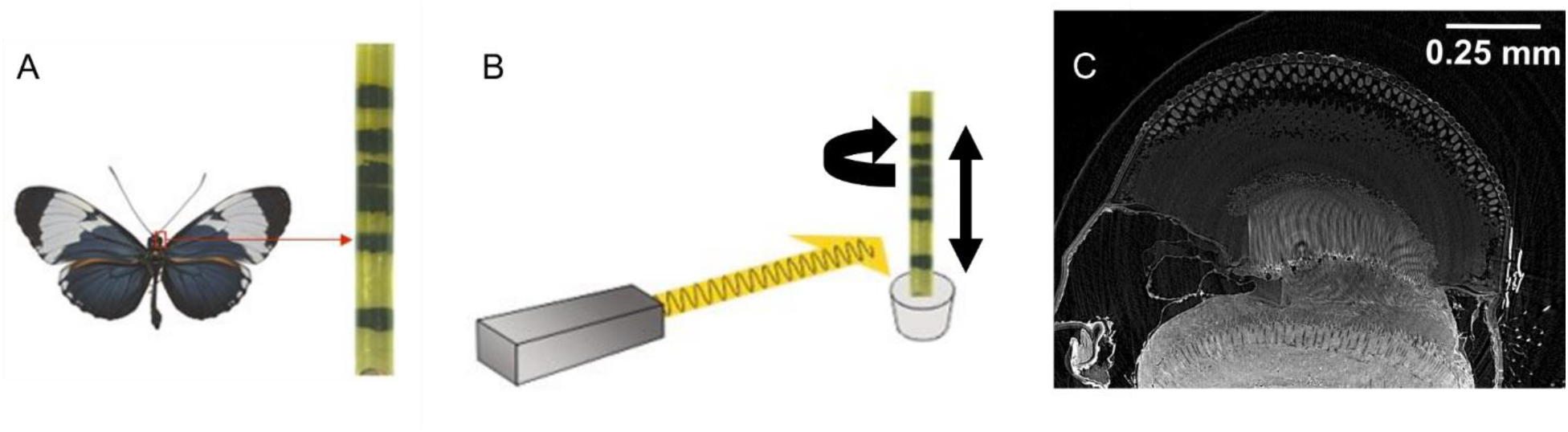
X-ray analysis pipeline showing A) diagram of insect eyes stacked in a vertical column, B) a diagram of the sample rotating and moving vertically in the X-ray beam, and C) a raw X-ray image. Butterfly in A) is from (Gallice, 2012).

We used an analysis pipeline to extract the relevant features from the X-ray datasets. For example, Fig 2A shows the segmentation output of ilastik (Berg et al., 2019), a free open-source software for image classification and segmentation. The output from ilastik gave us clusters of points corresponding to each crystalline cone, which we analyzed in Matlab (The Mathworks, Natick, MA) and Python (Van Rossum & Drake, 2009) (Fig, 2B, Fig. SI1).

**Figure 2.**
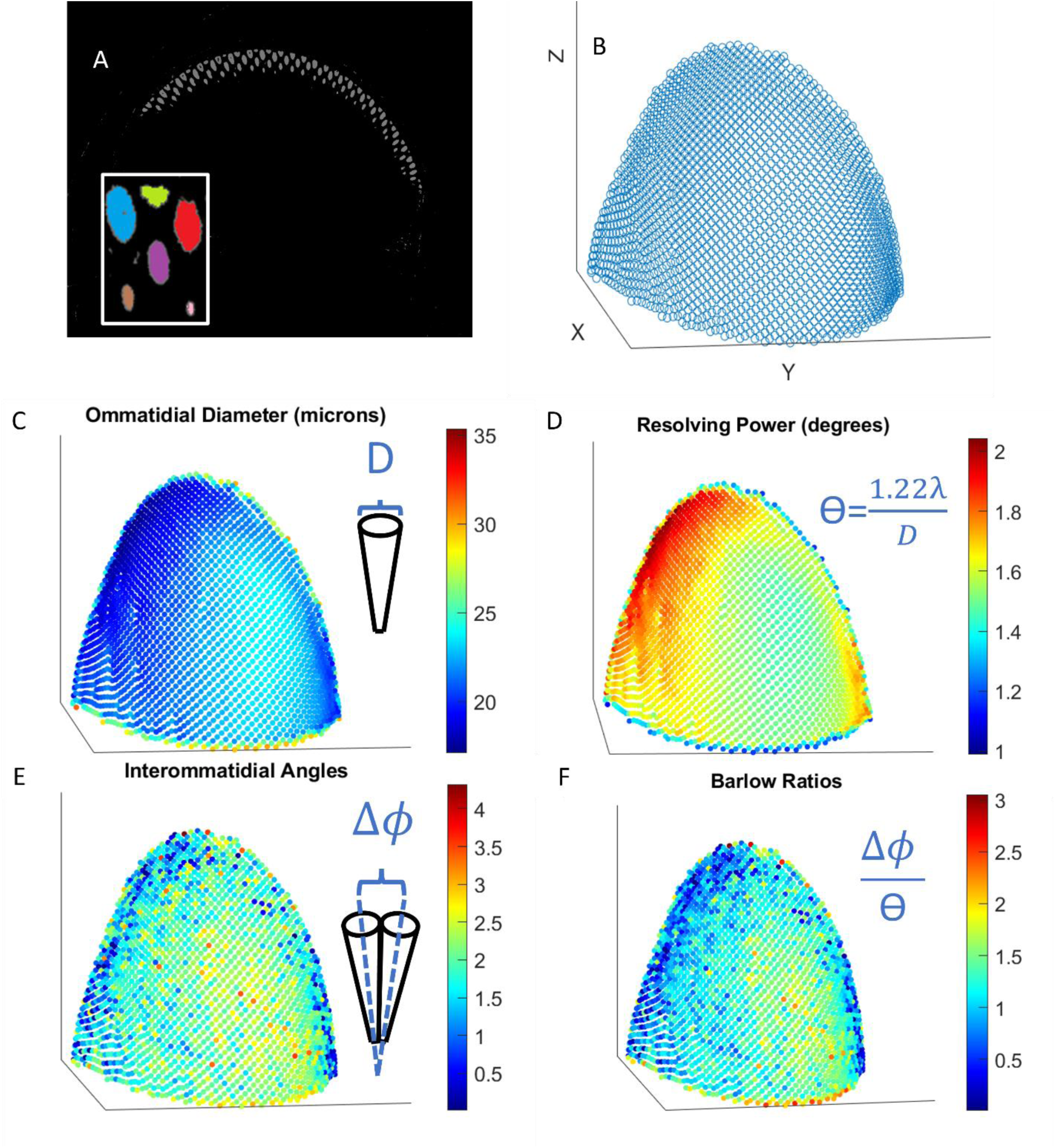
A) Segmented out crystalline cones with the inlay showing that cones are labeled as separate objects. B) Centers of crystalline cones plotted in Matlab where the rest of our analysis took place. C-F) show scatter plots showing how C) ommatidial diameter, D) resolving power, E) interommatidial angle, and F) Barlow ratios change across the eye. The portion of the eye shown here is from *Polites peckius*. Each point represents one ommatidium.

### Sample preparation

Insect samples were either collected in the wild in Chicago, IL (*Pieris rapae*, *Polites peckius*, and *Polygonia interrogationis*), collected from our breeding colonies at The University of Chicago (*Heliconius cydno*), or provided by Erica Westerman (University of Arkansas) (*Strymon melinus* and *Calycopis cecrops*). For dissections, insects were anesthetized by placing them at 4°C for ∼10 minutes. Insects were then submerged in ice cold Phosphate Buffered Saline (PBS) and dissected in PBS under a stereomicroscope to remove the cuticle outer layer and expose the brain. Brains with eyes intact were then cut from the body and submerged in fixative solution consisting of 0.1M Sodium Cacodylate buffer, pH 7.4, 2% paraformaldehyde, and 2.5% glutaraldehyde. Brains were incubated in fixative for ∼24hrs, gently rocking at 4°C. The next day, brains with eyes were prepared using electron microscopy protocols as previously described (Hua et al, 2015). Briefly, brains were washed extensively in cacodylate buffer at room temperature and stained sequentially with 2% osmium tetroxide (EMS) in cacodylate buffer, 2.5% potassium ferrocyanide (Sigma-Aldrich), thiocarbohydrazide, unbuffered 2% osmium tetroxide, 1% uranyl acetate, and 0.66% Aspartic acid buffered Lead (II) Nitrate with extensive rinses between each step with the exception of potassium ferrocyanide. The samples were then dehydrated in ethanol and propylene oxide and infiltrated with 812 Epon resin (EMS, Mixture: 49% Embed 812, 28% DDSA, 21% NMA, and 2.0% DMP 30). Samples were cured in custom cylindrical molds to stack multiple brains into one sample and to remove any edges to the resin that may affect X-ray imaging. The resin-infiltrated tissue was cured at 60°C for 3 days.

### µX-ray computed tomography

The syn-μCT data were acquired as previously described (Foxley et al, 2021). Briefly, we used the 32-ID beamline at the Advanced Photon Source, Argonne National Laboratory. The setup consists of a 1.8 cm-period undulator operated at a low deflection parameter value of *K* = 0.26. This yields a single quasi-monochromatic peak of energy 25 keV without the losses incurred by use of a <u>crystal monochromator</u>. For a sample 68 m from the undulator, this produces a photon fluence rate of about 1.8×107 photons s^-1^µm^-2^.

The x-rays were imaged using a 10 µm thick thin-film LuAG:Ce <u>scintillator</u> producing visible-light images then magnified using a 10X Mitutoyo long working distance microscope objective onto a 1920×1200 pixel CMOS camera (Point Gray GS3-U3-51S5M-C). The effective object space pixel size was 600 nm isotropic. The thickness of the thin-film scintillator matched the depth of focus of the objective lens, achieving a spatial resolution equivalent to the resolving power of the lens (1.3 µm for a NA of 0.21). Since the camera field of view was substantially smaller than the sample, a mosaic strategy was employed (Vescovi et al., 2018).

The sample was mounted on an air-bearing rotary stage (PI-Micos UPR-160 AIR) with motorized *x*/*y* translation stages located underneath and *x*/*y* piezo stages on top. Typical exposure time for a single projection image at one mosaic grid point and one rotation angle was 30 ms. 360° rotation angles were used at each grid point. The sample was translated through a 6 ×18 tomosaic grid.

#### Data Analysis

Crystalline cones from the raw x-ray datasets were segmented using the software ilastik and code based off cc3d (Silversmith, 2021). This generated sets of voxels corresponding to each of the crystalline cones. Outliers in the set of points that were not part of the cones were deleted manually.

We defined the center of each crystalline cone as its center of mass. Then we estimated the local radius of the eye by fitting a sphere to clusters of 60 points corresponding to the crystalline cone centers. We chose 60 points because this encompasses a hexagonal array surrounding a single point extending 4 ommatidia out in all directions. Vectors from the center of the sphere to the center of each cone were calculated. Once we have defined the ‘center’ of the eye from the local curvature we can then use the vectors from that putative center to the centers of the cones to define an ommatidial angle. The angles between a cone’s vector and its six nearest neighbors’ vectors were averaged, and this was used as the (local) interommatidial angle (Δɸ). The average distance to the six nearest neighbors was used as the diameter of the ommatidium (D). Since the center of each cone lies below the surface of each eye facet, this systematically underestimates the value of D by potentially a significant fraction of the cone length times Δɸ (measured in radians). This systematic error is then of order 2 microns or less, which is considerably smaller than both the mean and the variance of D (Table SI1). Resolving power was calculated by θ=1.22*λ/D where λ is the wavelength of light. For our analysis, λ=500 nm, as it corresponds to broad peaks in both the typical sunlight spectrum and photoreceptor sensitivity in many insects. This is also the λ that Barlow used for his calculations. The ratio Δɸ/θ, aka the Barlow ratio, was also calculated. Extreme outliers were cut off when we noted corresponding defects in the x-ray images or where the values seemed biologically implausible (e.g., interommatidial angles greater than 90 degrees). These outliers occurred almost exclusively at the edges of the eyes. Cone shapes were determined by centering and overlaying all cones within a single eye, and keeping the collection of shared points, points with at least 25% overlap, across all cones. This was done to reduce noise in the segmentation of individual cones. The number of cones overlayed per eye varied from about 600 to 3000. There appeared to be some variation in cones across the eye, but the biggest deviations from the average seemed to come from the cones at the outer edges of the eye. Then, the boundary of the cone was calculated from this set of overlapping points using the “boundary” function on Matlab. We used the Pearson correlation coefficient to determine the relationship between wingspan and cone length as well as wingspan and cone ratio. We also calculated the aspect ratio and cone ratio for each individual cone.

## Results

In order to understand whether ommatidial diameter, interommatidial angles, resolving power, and Barlow ratios changed within individuals, we first looked at how these different parameters varied across the surface of a single eye. When creating these maps, we found that ommatidial diameter (as well as resolving power) appeared to vary gradually from areas with larger diameters up to 35.3 microns to those with smaller diameters down to 17.1 microns (Fig 2C, D). In contrast, the changes in interommatidial angle and Barlow ratio across the eye were not as smooth, with transitions from higher to lower acuity areas being less clear by visual inspection (Fig 2E, F).

Next, we asked how these parameters varied across different individuals by reporting the statistics of the distributions for these variables for each individual. The average median ommatidial diameter across all individuals was 24.5 microns, ranging from 20.54 to 31.09 microns, with an average interquartile range of 3.36 (Fig 3A). Resolving powers had an average median value of 1.46 degrees, ranging from 1.12 to 1.70 degrees, with an average interquartile range of 0.18 (Fig 3B). The median interommatidial angle measured across all species ranged from 1.42 to 1.87 degrees, with an average of 1.66 degrees and an average interquartile range of 1.01 degrees (Fig 3B). Medians for the Barlow ratio ranged from 0.846 to 1.49 with two individuals having medians within the optimal range. The average median Barlow ratio was 1.17 with an average interquartile range of 0.80 (Fig 4A).

**Figure 3.**
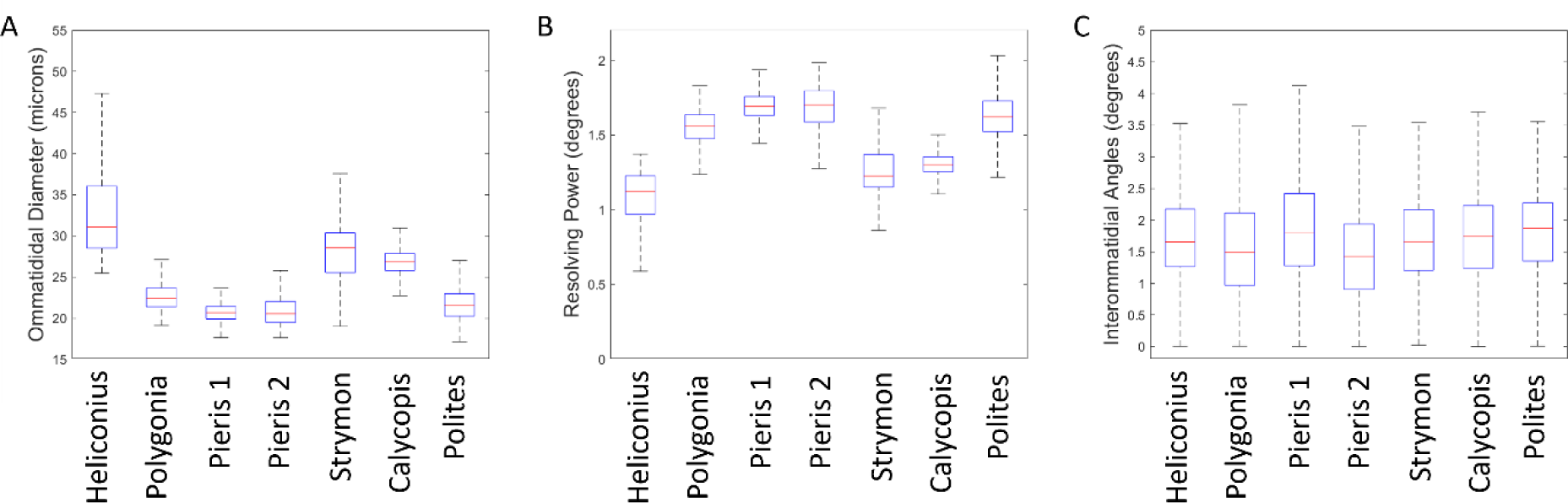
Plots showing the median (red line) A) ommatidial diameter in microns, B) resolving powers, C) and interommatidial angles in degrees for the seven individuals. Boxes show interquartile range and whiskers show the lower and upper quartiles (Outliers are shown in Fig. SI2 for clarity of presentation).

**Figure 4.**
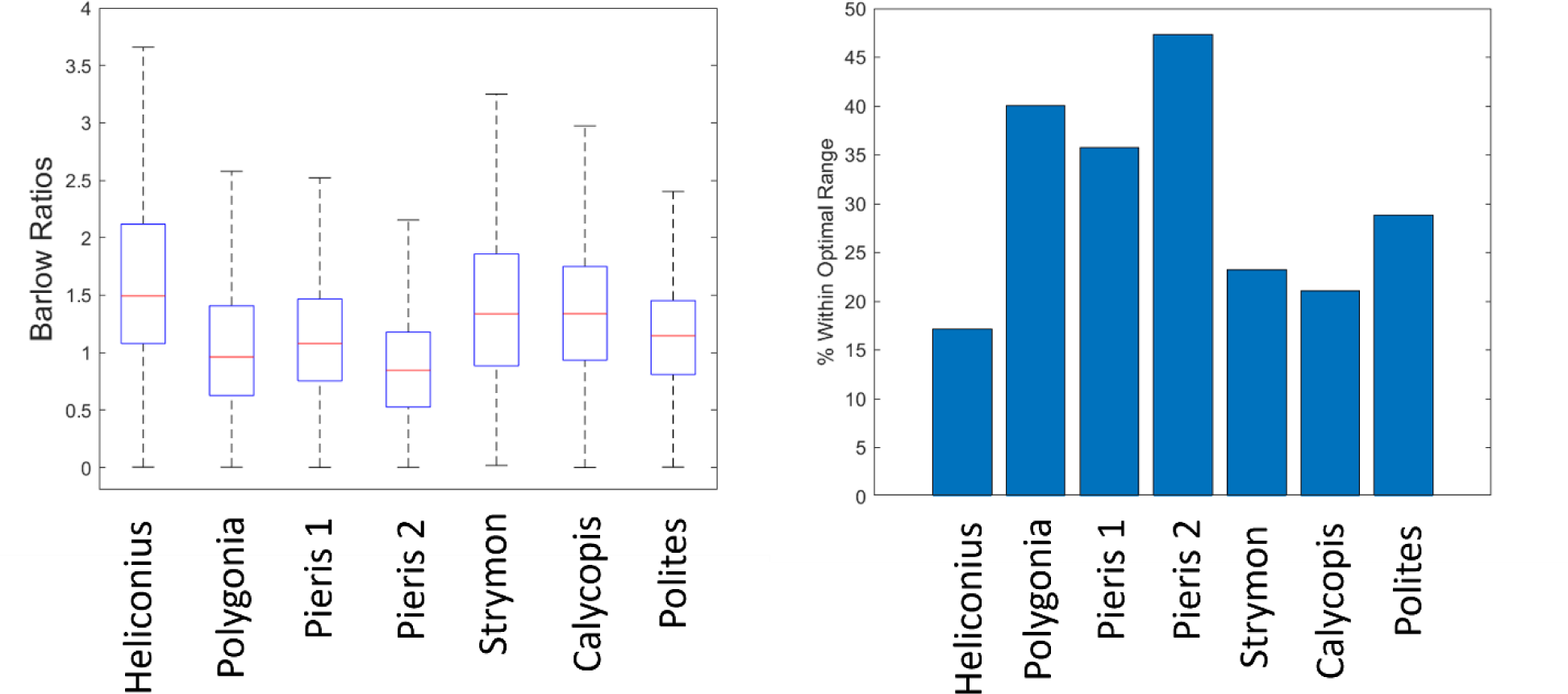
A) Boxplot showing the median (red line) Barlow ratio for the seven individuals. Boxes show interquartile range and whiskers show the lower and upper quartiles (Outliers are shown in Fig. SI2). B) Bar plot showing the percentage of each portion of eye with a Barlow ratio that fell between 0.4 and 1.

Because our method provided great enough resolution to clearly distinguish whole crystalline cones, we decided to look at the micron scale morphology of this structure, which guides the light focused by the cornea and lens to the ommatidia. When looking at the size and shape of this structure, we found variation across individuals that would need to be disentangled from variation across the eye with larger datasets. For instance, the average “typical” cone length across species (Fig. 5) was 44.4 microns, but ranged from 22.2 microns to 72.0 microns, and some cones had a defined point at the bottom while others were more rounded. In order to quantify how tapered a cone was, we took the ratio of the cone diameter at 10% of the length from the bottom and the maximum diameter at the top and found the mean of this ratio was 0.3669. The most tapered cone had a ratio of 0.2121, while the least tapered cone was less than half as tapered with a ratio of 0.5362. When analyzing morphological data, it is important to consider scaling relationships in the data (Jablonski et al, 1996). In our data, when examining allometric relationships, we found that there was a negative correlation between the wingspan of a species and the typical cone length (r(7)=-0.8881, p=0.0076) as well as a negative correlation between wingspan and cone ratio (r(7)=-0.7949, p=0.0326). Besides looking at the typical crystalline cones, we also looked at the shapes of all the cones across each eye. Changes in different parameters such as the length (Fig 5B), aspect ratio (Fig 5C), and cone ratio (Fig 5D) could be seen across the eyes of all individuals.

**Figure 5.**
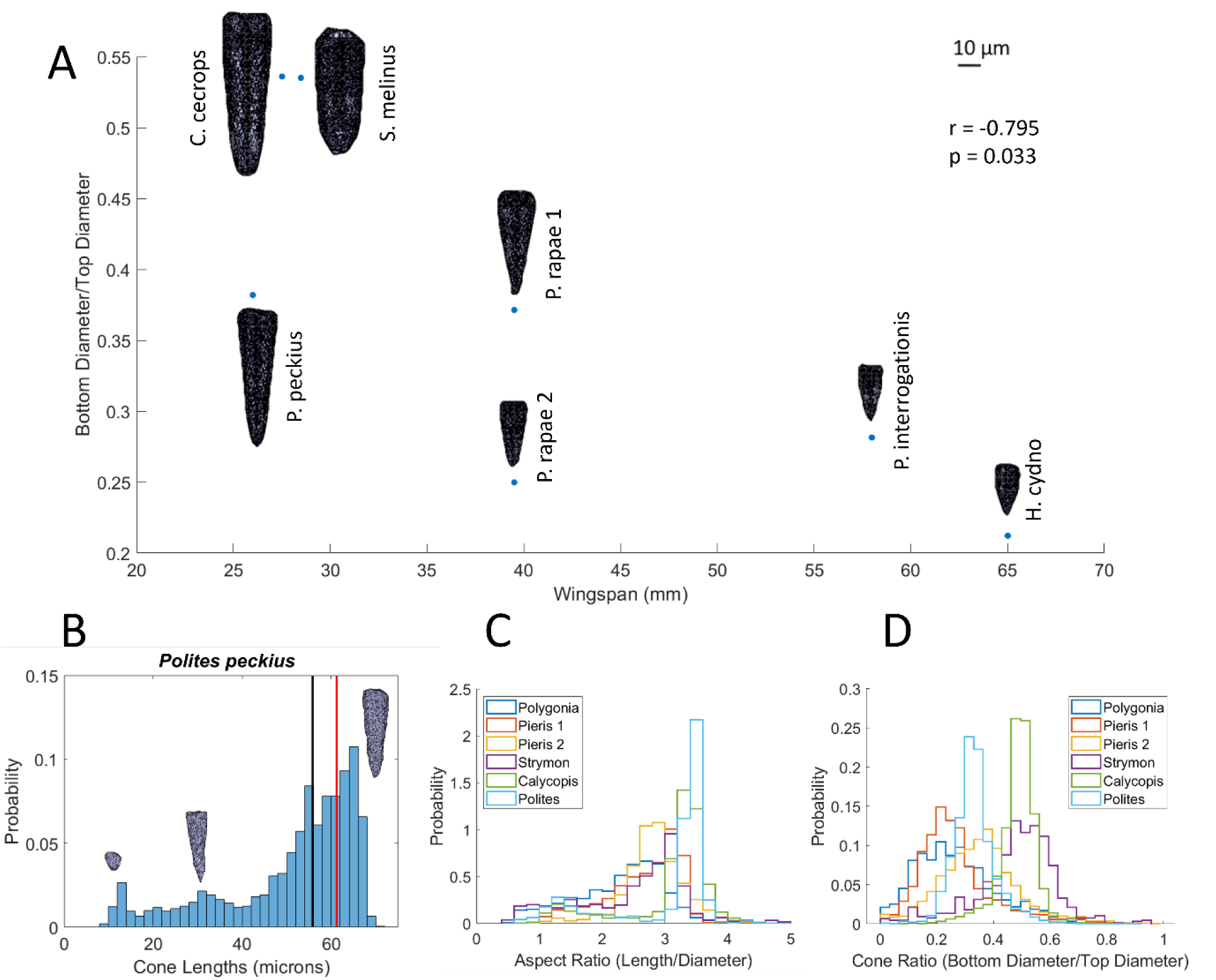
A) Cone shapes from all 7 individuals plotted by the length of the cone and the ratio of the diameter of the cone 10% from the bottom and the maximum width of the cone at the top. Scale bar shows 10 microns. Black line denotes the median cone length, while the red line shows the cone length of the typical cone. B) Histogram showing the cone lengths across the eye of Polites peckius with example cones from the 3 different peaks in the distribution. C) Histograms showing the aspect ratios and D) cone ratios of each individual (excluding Heliconius as there were issues with individual cone segmentation in this dataset).

## Discussion

Our method provides a new way to study insect morphology, especially the individual components of the eye, using a higher contrast staining method, higher resolution syn-μCT, and a novel analysis pipeline. With the method, an entire eye can be surveyed for microscopic features like ommatidial diameters, angles, and cone morphologies. Previous methods measuring microscopic features have either imaged smaller volumes at higher resolution (e.g. Hao et al., 2023) or larger volumes at lower resolution (e.g. Currea et al., 2023). Using our approach, we were able to analyze seven insects of six different butterfly species to show that the Barlow ratio of the ommatidia falls in or near the theoretical optimum, but notable portions of the eye have Barlow ratios greater than this optimum. This suggests that portions of the visual scene are undersampled. In previous work that has shown a similar kind of spatial undersampling in insect eyes, it has been suggested that this is due to motion blur from the animal moving about its environment (Land, 1997). This indicates that theoretical models must also account for the angular velocity of the organism or objects in its visual environment and other processing that happens later in the visual system to optimize an insect’s vision. Additional research is needed to assess how different luminance may change the Barlow ratio and if the theoretical models that account for light levels, as described by Snyder et al (1977), are correct.

### Limitations

There are several limitations to this study. One issue is that our imaging often did not cover the entire eye. While this is not ideal, we were still able to analyze large enough portions of the eyes to capture the variability across eyes. This study is complementary to previous work that allows sampling of different parts of the eye. However, there is a future planned upgrade to the APS synchrotron that will enable imaging of entire eyes and nervous systems of insects (Argonne National Laboratory, n.d.).

Another limitation in study is that sample preparation for electron microscopy is well known to change the native structures of brain tissue (Zhang et al., 2017). However, most of these artifacts involve changes in the volume of the extracellular space (Van Harreveld & Steiner, 1970; Pallotto et al., 2015). We analyzed crystalline cones, which are composed of concentrated, hydrophobic proteins in closely related moths and likely less susceptible to dehydration-based distortions (Schlamp, 1989). We designed an analysis pipeline robust to small changes in orientation, thereby preserving local curvature and diameters.

Finally, we see smooth variation across individual eyes, which gives us confidence that the differences observed are not simply noise from artifacts.

### Comparison to prior work

Previous analyses of ommatidial diameter and interommatidial angles were done using light microscopy (and more recently fluorescence microscopy) and analyzed manually (Horridge, 1978; Rutowski & Warrant, 2002; Baumgartner, 1928; del Portillo, 1936; Rigossi et al 2021), and therefore would take a much longer time to collect data on the same volume of eye. Previous methods for calculating interommatidial angles include observing how many ommatidia pseudopupils crossed while rotating the eye a certain angle, using the optomotor response, and manually measuring histological sections (Horridge, 1978; Rutowski & Warrant, 2002; Gotz, 1965; Baumgartner, 1928; del Portillo, 1936; Rigossi et al., 2021). All of these methods are subject to human error, but our method provides an automated way to calculate both the interommatidial angle and the ommatidial diameter.

Several newer methods have been proposed for measuring interommatidial angles and other eye parameters. One such method involves staining photoreceptors with fluorescent dyes to measure interommatidial angles using the pseudopupil in insects with dark eyes (Rigossi et al., 2021). One advantage of this fluorescence method is that it can be done using live animals and avoids any distortion that may occur during sample preparation. However, this is the only parameter that can be measured with this technique and cannot reveal the morphology of internal structures.

The µCT method has recently been used to measure angles and other eye parameters in bees (Taylor et al, 2019) and ommatidial diameters in other compound eyes (Currea et al., 2023), but our method using the 32-ID beamline achieves ∼18x or ∼170x greater resolution respectively, and imaging speeds of ∼1 mm^3^/30min. This enhanced resolution combined with our novel embedding method allows for greater automated throughput. Furthermore, conventional lab-based µCT imaging that can achieve comparable spatial resolution (Alba-Tercedor et al., 2021) have worse contrast resolution than syn-µCT (Goyens et al., 2018). Additionally, our staining method provides even greater contrast and allows us to better see crystalline cones, whereas previous µCT reconstructions were unable to capture this structure. We found large variation in the sizes and shapes of the typical crystalline cones across individuals and especially species. Using our 3D models of these cones, further research can be done to explore how light passes through these structures and impinges on the rhabdom.

### The Barlow Ratio, Variation, and Motion Blur

There was considerable variation in all measurements across individual eyes. All the species had average Barlow ratios near the theoretical optimum, but large portions of each eye had ratios that were greater than expected, meaning the interommatidial angle was greater than the resolving power, suggesting the visual scene is undersampled. Undersampling of the visual scene has been observed in other insects. In previous studies that have measured a similar ratio, the acceptance angle to the interommatidial angle, in other diurnal insects also found that the visual scene was undersampled (Land 1997). One reason insect vision might be undersampled is to account for motion blur. For example, according to a paper from Snyder et al, the fly *Musca* would have a Barlow ratio of 2.13, which is greater than both Barlow’s optimum and the optimum calculated by Snyder et al (1977). However, this value did approach a value that Snyder *et al*. deemed more reasonable once angular velocity of the insect was accounted for. Finally, Snyder *et al*. also looked at how different light levels would affect the theoretical optimum for p=D*Δɸ=0.61*(Δɸ/θ). They theorized that in lower light conditions, the optimal p would be larger. Further research must be done to examine crepuscular and nocturnal Lepidoptera to determine if this is indeed the case.

Finally, we determined the morphology of a typical crystalline cone for each species. We found considerable variation in height and width within and across individuals, which could be due to scaling with total body size or cone density within the eye. Previous analyses have identified that the point-like end of the crystalline cone corresponds with the focal point of the lens, allowing the most efficient transfer of photons into a single rhabdom (Schwarz et al., 2011). For species adapted to low light, the hypothesis is instead that cones are larger and more bulbous with the focal point well inside the cone, which is believed to confer an advantage for greater light collection by transmission through a ‘clear zone’ to multiple rhabdoms (Warrant, 2017). Since the crystalline cone’s function is to funnel light onto the rhabdom, further studies could potentially determine how cone optics vary across the eye and between species. Measurements of body size that incorporate forewing length are correlated with larger eye sizes (Seymoure et al., 2015), and longer cones correspond to smaller wingspans, suggesting smaller Lepidoptera have flatter lenses as they have a longer focal length. Syn-µCT also clearly shows the shape of the lens, so our method would be useful in testing this hypothesis. Future work will explore these differences more fully, by modeling the wave optics of light passing through lenses and cones with these different shapes.

## Conflict of Interest Statement

We have no conflicts of interest to disclose.

## Author Contributions

Gregg A. Wildenberg, Narayanan Kasthuri, Stephanie E. Palmer, and Dawn Paukner conceived the ideas and designed methodology; Gregg Wildenberg collected the data; Dawn Paukner and Griffin S. Badalamente created the analysis pipeline and analyzed the data; Dawn Paukner and Narayanan Kasthuri led the writing of the manuscript. Dawn Paukner, Gregg A. Wildenberg, Peter B. Littlewood, Marcus R. Kronforst, Stephanie E. Palmer, and Narayanan Kasthuri contributed to the interpretation of the results. All authors contributed critically to the drafts and gave final approval for publication.

## Data Availability

Raw X-ray datasets will be made available on BossDB. Code will be made available on github.com/dpaukner.

## Supplementary Figures

**Figure SI1.**
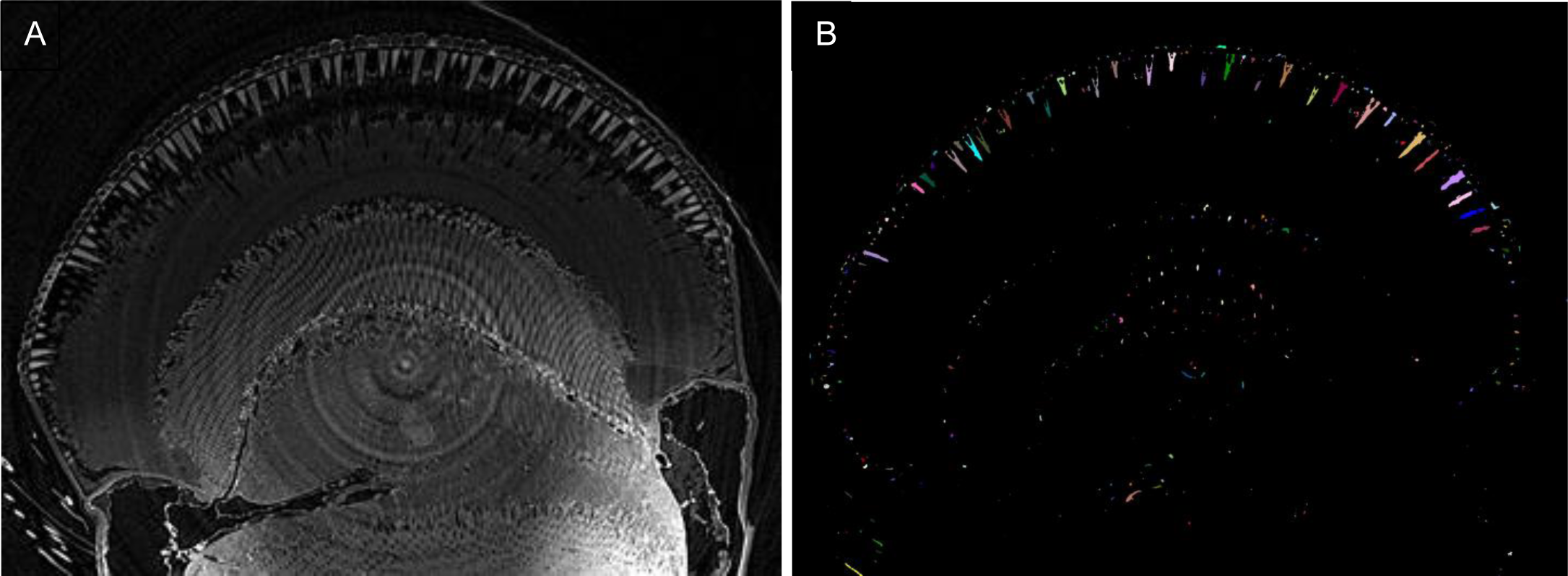
A) Gif moving through the raw x-ray stack of the *Polites peckius* eye. B) Gif moving through a stack of the *Polites peckius* eye after segmentation of crystalline cones.

**Figure SI2.**
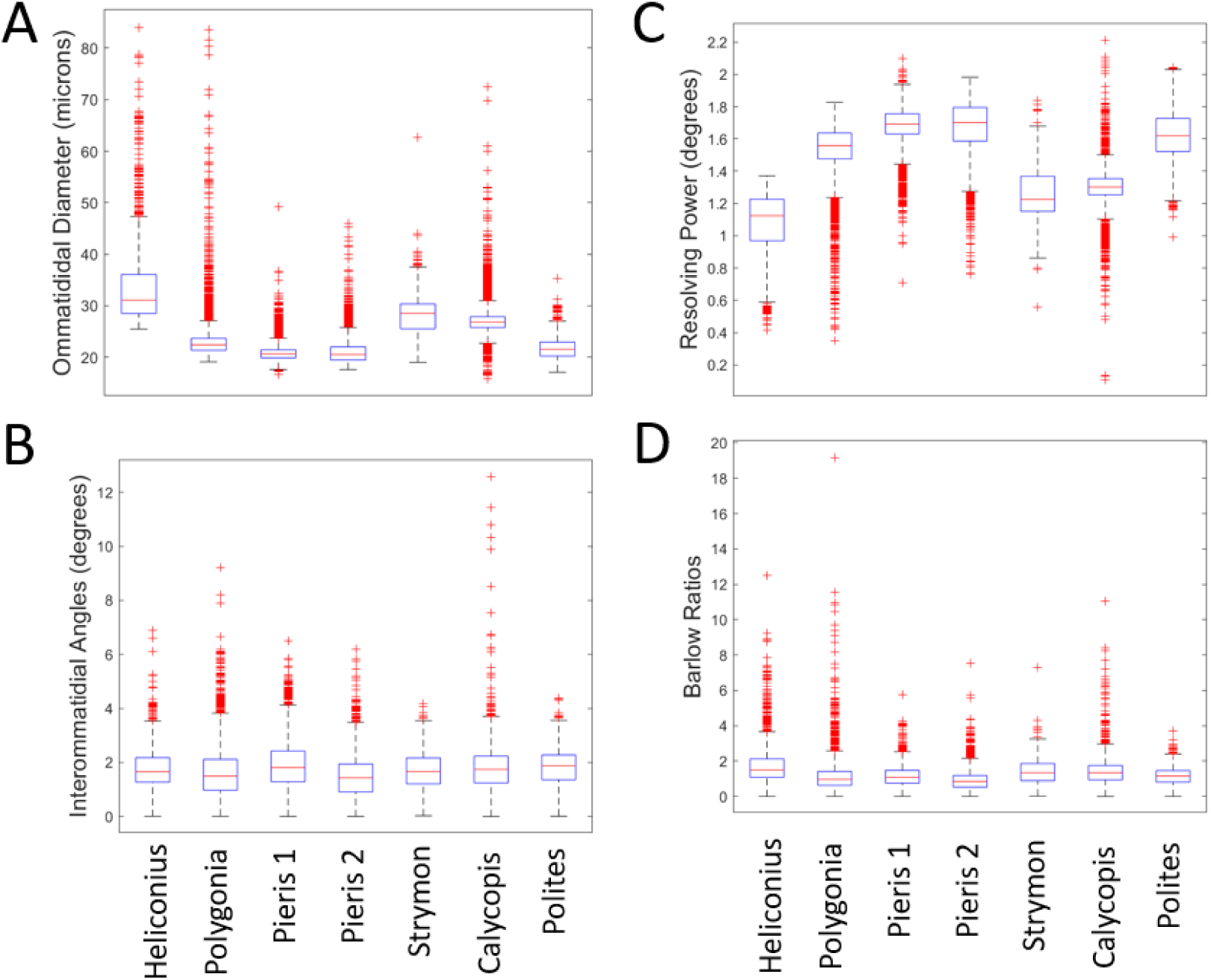
Boxplots showing A) ommatidial diameters, B) interommatidial angles, C) resolving powers, and D) Barlow ratios. Red lines show medians, boxes show interquartile ranges, and whiskers show the lower and upper quartiles. Red plus signs denote outliers.

**Table SI1.** Shows the estimated error, mean, and variance of ommatidial diameters for each indiviudal.

|  | *H.c. galanthus* | *P. interrogationis* | *P. rapae 1* | *P. rapae 2* | *S. melinus* | *C. cecrops* | *P. peckius* |
| --- | --- | --- | --- | --- | --- | --- | --- |
| Estimated Error | 0.684722884 | 0.725894193 | 1.511149047 | 0.743897027 | 1.628931642 | 2.230212656 | 1.924340469 |
| Mean D | 33.75005873 | 23.74537682 | 21.03077 | 21.18907642 | 28.52390497 | 27.39373852 | 21.6664377 |
| Variance of D | 60.89412798 | 30.30223407 | 4.000139676 | 6.904767902 | 16.57940237 | 16.12510145 | 4.460714212 |

## Notes

### Competing Interest Statement

The authors have declared no competing interest.

## References

1. Alba-Tercedor, J., Hunter, W.B. & Alba-Alejandre, I. Using micro-computed tomography to reveal the anatomy of adult Diaphorina citri Kuwayama (Insecta: Hemiptera, Liviidae) and how it pierces and feeds within a citrus leaf. Sci Rep 11, 1358 (2021). 10.1038/s41598-020-80404-z

2. Argonne National Laboratory. (n.d.). APS Upgrade. https://www.aps.anl.gov/APS-Upgrade

3. Barlow, H. B. (1952). The Size of Ommatidia in Apposition Eyes. The Journal of Experimental Biology, 25(*4*), 667–674.

4. Baumgärtner, H. (1928). Der formensinn und die sehschärfe der bienen. Zeitschrift für vergleichende Physiologie, 7(1), 56–143.

5. Baxter, S. W., Nadeau, N. J., Maroja, L. S., Wilkinson, P., Counterman, B. A., Dawson, A., … & Jiggins, C. D. (2010). Genomic hotspots for adaptation: the population genetics of Müllerian mimicry in the Heliconius melpomene clade. PLoS genetics, 6(2), e1000794.

6. Berg, S., Kutra, D., Kroeger, T. et al. ilastik: interactive machine learning for (bio)image analysis. Nat Methods 16, 1226–1232 (2019). 10.1038/s41592-019-0582-9

7. Butler, P. M. (1963). Tooth morphology and primate evolution. Dental anthropology, 1–13.

8. Chown, S. L., & Terblanche, J. S. (2006). Physiological diversity in insects: ecological and evolutionary contexts. Advances in insect physiology, 33, 50–152.

9. Currea, J. P., Sondhi, Y., Kawahara, A. Y., & Theobald, J. (2023). Measuring compound eye optics with microscope and microCT images. Communications Biology, 6(1), 246.

10. Darwin, C. (1859). The Origin of Species by Means of Natural Selection, Or, The Preservation of Favoured Races in the Struggle for Life. Books, Incorporated, Pub..

11. del Portillo, J. (1936). Beziehungen zwischen den Öffnungswinkeln der Ommatidien, Krümmung und Gestalt der Insektenaugen und ihrer funktionellen Aufgabe. Zeitschrift für vergleichende Physiologie, 23(1), 100–145.

12. Dobzhansky, T. (1982). Genetics and the Origin of Species (No. 11). Columbia university press.

13. Dyer, E. L., Gray Roncal, W., Prasad, J. A., Fernandes, H. L., Gürsoy, D., De Andrade, V., Fezzaa, K., Xiao, X., Vogelstein, J. T., Jacobsen, C., Körding, K. P., & Kasthuri, N. (2017). Quantifying Mesoscale Neuroanatomy Using X-Ray Microtomography. eNeuro, 4(5), ENEURO.0195-17.2017. 10.1523/ENEURO.0195-17.2017

14. Foxley, S., Sampathkumar, V., De Andrade, V., Trinkle, S., Sorokina, A., Norwood, K., La Riviere, P., & Kasthuri, N. (2021). Multi-modal imaging of a single mouse brain over five orders of magnitude of resolution. NeuroImage, 238, 118250. 10.1016/j.neuroimage.2021.118250

15. Friedrich, F., Matsumura, Y., Pohl, H., Bai, M., Hörnschemeyer, T., & Beutel, R. G. (2014). Insect morphology in the age of phylogenomics: innovative techniques and its future role in systematics. Entomological Science, 17(1), 1–24.

16. Gallice, G (2012). Heliconius cydno galanthus [Photograph]. https://www.flickr.com/photos/dejeuxx/6626128815/in/photostream.

17. Götz, K.G. Die optischen Übertragungseigenschaften der Komplexaugen von Drosophila. Kybernetik 2, 215–221 (1965). 10.1007/BF00306417

18. Goyens, J., Vasilopoulou-Kampitsi, M., Claes, R., Sijbers, J. and Mancini, L. (2018), Enhanced contrast in X-ray microtomographic images of the membranous labyrinth using different X-ray sources and scanning modes. J. Anat., 233: 770–782. 10.1111/joa.12885

19. Hao, Y., Wang, Q., Wen, C., & Wen, J. (2023). Comparison of Fine Structure of the Compound Eyes in Eucryptorrhynchus scrobiculatus and Eucryptorrhynchus brandti Adults. Insects, 14(8), 699. 10.3390/insects14080699

20. Horridge G. A. (1978). The separation of visual axes in apposition compound eyes. Philosophical transactions of the Royal Society of London. Series B, Biological sciences, 285(1003), 1–59. 10.1098/rstb.1978.0093

21. Hua, Y., Laserstein, P., & Helmstaedter, M. (2015). Large-volume en-bloc staining for electron microscopy-based connectomics. Nature communications, 6, 7923. 10.1038/ncomms8923

22. Jablonski, D., Erwin, D. H., & Lipps, J. H. (1996). Body size and macroevolution. Evolutionary paleobiology. University of Chicago Press, Chicago, 256–289.

23. Johnson, J. T., Hansen, M. S., Wu, I., Healy, L. J., Johnson, C. R., Jones, G. M., … & Keller, C. (2006). Virtual histology of transgenic mouse embryos for high-throughput phenotyping. PLoS Genetics, 2(4), e61.

24. Kronforst, M. R., Young, L. G., Kapan, D. D., McNeely, C., O’Neill, R. J., & Gilbert, L. E. (2006). Linkage of butterfly mate preference and wing color preference cue at the genomic location of wingless. Proceedings of the National Academy of Sciences, 103(17), 6575–6580.

25. Land M. F. (1997). Visual acuity in insects. Annual review of entomology, 42, 147–177. 10.1146/annurev.ento.42.1.147

26. MATLAB. (2021). version 9.10.0.1669831 (R2021a) Update 2. Natick, Massachusetts: The MathWorks Inc.

27. Namiki, S., & Kanzaki, R. (2018). Morphology of visual projection neurons supplying premotor area in the brain of the silkmoth Bombyx mori. Cell and tissue research, 374(3), 497–515. 10.1007/s00441-018-2892-0

28. Pallotto, M., Watkins, P. V., Fubara, B., Singer, J. H., & Briggman, K. L. (2015). Extracellular space preservation aids the connectomic analysis of neural circuits. Elife, 4, e08206.

29. Poulton, E. B. (1909). Mimicry in the butterflies of North America. Annals of the entomological Society of America, 2(4), 203–242.

30. Price, P. W., Denno, R. F., Eubanks, M. D., Finke, D. L., & Kaplan, I. (2011). Insect ecology: behavior, populations and communities. Cambridge University Press.

31. Ribi, W., Senden, T. J., Sakellariou, A., Limaye, A., & Zhang, S. (2008). Imaging honey bee brain anatomy with micro-X-ray-computed tomography. Journal of neuroscience methods, 171(1), 93–97.

32. Rigosi, E., Warrant, E. J., & O’Carroll, D. C. (2021). A new, fluorescence-based method for visualizing the pseudopupil and assessing optical acuity in the dark compound eyes of honeybees and other insects. Scientific reports, 11(1), 21267. 10.1038/s41598-021-00407-2

33. Rutowski, R.L., Warrant, E.J. Visual field structure in the Empress Leilia, *Asterocampa leilia* (Lepidoptera, Nymphalidae): dimensions and regional variation in acuity. J Comp Physiol A 188, 1–12 (2002). 10.1007/s00359-001-0273-7

34. Schlamp, Cassandra Lee, “Composition, Structure And Development Of The Crystalline Cone Of The Superposition Compound Eye” (1989). Digitized Theses. 1851. https://ir.lib.uwo.ca/digitizedtheses/1851

35. Schwarz, S., Narendra, A., & Zeil, J. (2011). The properties of the visual system in the Australian desert ant Melophorus bagoti. Arthropod structure & development, 40(2), 128–134. 10.1016/j.asd.2010.10.003

36. Seymoure, B. M., Mcmillan, W. O., & Rutowski, R. (2015). Peripheral eye dimensions in Longwing (Heliconius) butterflies vary with body size and sex but not light environment nor mimicry ring. J Res Lepid, 48, 83–92.

37. Silversmith, W. (2021). cc3d: Connected components on multilabel 3D & 2D images. (Version 3.2.1) [Computer software]. https://zenodo.org/record/5535251

38. Snyder, A.W., Stavenga, D.G. & Laughlin, S.B. Spatial information capacity of compound eyes. J. Comp. Physiol. 116, 183–207 (1977). 10.1007/BF00605402

39. Stork N. E. (2018). How Many Species of Insects and Other Terrestrial Arthropods Are There on Earth?. Annual review of entomology, 63, 31–45. 10.1146/annurev-ento-020117-043348

40. Taylor, G. J., Tichit, P., Schmidt, M. D., Bodey, A. J., Rau, C., & Baird, E. (2019). Bumblebee visual allometry results in locally improved resolution and globally improved sensitivity. eLife, 8, e40613. 10.7554/eLife.40613

41. Van de Kamp, T., Rolo, T.O.D & Baumbach, T. (2014). Scanning the past– synchrotron X-ray microtomography of fossil wasps in amber. Entomologie heute, 26, 151–160.

42. Van den Boogert, T., Van Hoof, M., Handschuh, S., Glueckert, R., Guinand, N., Guyot, J. P., … & Van de Berg, R. (2018). Optimization of 3D-visualization of micro-anatomical structures of the human inner ear in osmium tetroxide contrast enhanced micro-CT scans. Frontiers in Neuroanatomy, 12, 41.

43. Van Harreveld, A., & Steiner, J. (1970). Extracellular space in frozen and ethanol substituted central nervous tissue. The Anatomical Record, 166(1), 117–129.

44. Van Rossum, G., & Drake, F. L. (2009). Python 3 Reference Manual. Scotts Valley, CA: CreateSpace.

45. Vescovi, R., Du, M., Andrade, V. D., Scullin, W., Gürsoy, D., & Jacobsen, C. (2018). Tomosaic: efficient acquisition and reconstruction of teravoxel tomography data using limited-size synchrotron X-ray beams. Journal of synchrotron radiation, 25(5), 1478–1489.

46. Warrant E. J. (2017). The remarkable visual capacities of nocturnal insects: vision at the limits with small eyes and tiny brains. *Philosophical transactions of the Royal Society of London. Series B*, Biological sciences, 372(1717), 20160063. 10.1098/rstb.2016.0063

47. Wipfler, B., Pohl, H., Yavorskaya, M. I., & Beutel, R. G. (2016). A review of methods for analysing insect structures—the role of morphology in the age of phylogenomics. Current opinion in insect science, 18, 60–68.

48. Zhang, Y., Huang, T., Jorgens, D. M., Nickerson, A., Lin, L. J., Pelz, J., Gray, J. W., López, C. S., & Nan, X. (2017). Quantitating morphological changes in biological samples during scanning electron microscopy sample preparation with correlative super-resolution microscopy. PloS one, 12(5), e0176839. 10.1371/journal.pone.0176839

