## Supplementary figures and images for "Synchrotron-source micro-x-ray computed tomography for examining butterfly eyes"

### Supplemental Figure 1B

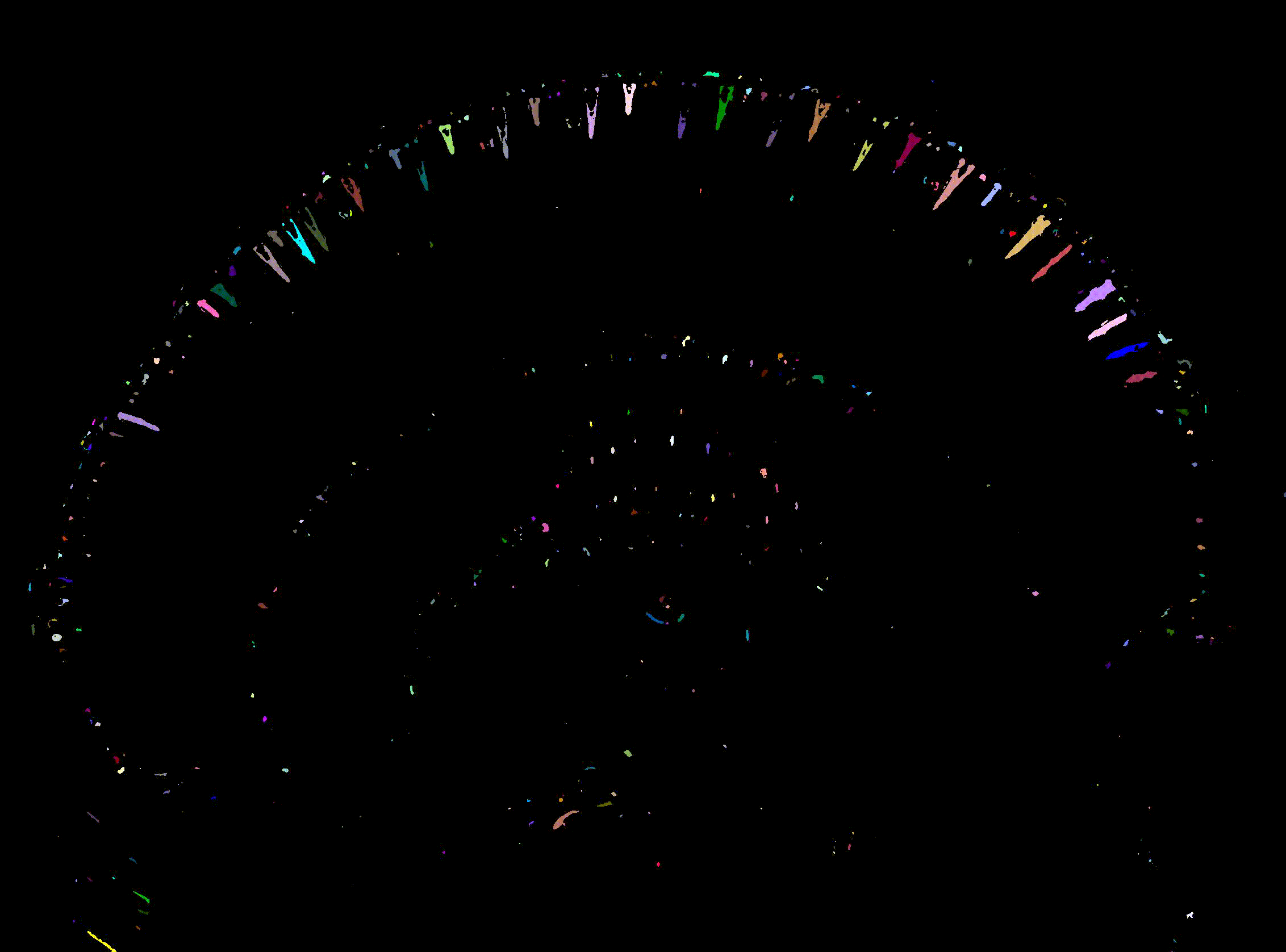
